# Generalise or Memorise? Benchmarking Ligand-Conditioned Protein Generation from Sequence-Only Data

**DOI:** 10.64898/2026.02.06.704305

**Authors:** Alex Vicente-Sola, Lars Dornfeld, Joan Coines, Noelia Ferruz

## Abstract

Proteins can bind small molecules with high specificity. However, designing proteins that bind userdefined ligands remains a challenge, typically relying on structural information and costly experimental iteration. While protein language models (pLMs) have shown promise for unconditional generation and conditioning on coarse functional labels, instance-level conditioning on a specific ligand has not been evaluated using purely textual inputs. Here we frame small-molecule protein binder design as a sequence-to-sequence translation problem and train ligand-conditioned pLMs that map molecular strings to candidate binder sequences. We curate large-scale ligand–protein datasets (>17M ligand-protein pairs) covering different data regimes and train a suite of models, spanning 16 to 700M parameters. Results reveal a consistent trade-off driven by supervision ambiguity: when each ligand is paired with few proteins, models generate near-neighbour, foldable sequences; when each ligand is paired with many proteins, generations are more diverse but less consistently foldable. Our study exposes how annotation diversity and sampling choices elicit this behaviour and how it changes with the data distribution. These insights highlight dataset redundancy and incompleteness as key bottlenecks for sequence-only binder design. We release the curated datasets, trained models, and evaluation tools to support future work on ligand-conditioned protein generation.

## 1. Introduction

Proteins can sense small molecules with exceptional sensitivity, initiating signaling cascades, transferring information, modulating the activity of other proteins, or catalyzing reactions that generate bioactive compounds and polymers. This functional diversity has long fascinated scientists and inspired efforts to engineer artificial proteins that recognize user-defined ligands. Yet, despite major progress, designing small-molecule binders remains exceptionally difficult: the protein must fold correctly and present side chains in a precise geometry that allows ligand access and stable binding, with even subtle deviations often abolishing affinity. Consequently, most successful strategies begin with weakly binding scaffolds and enhance performance through laboratory-directed evolution, aside from a few notable exceptions(Kortemme, 2024).

In recent years, protein design has been transformed by methods that leverage large-scale data and machine learning (ML) architectures. In this context, protein language models (pLMs) have shown strong performance in generating proteins that explore sequence space while retaining natural-like properties, all without the need for structural annotation. These advances have enabled the design of diverse proteins and enzymes, including lysozymes(Madani et al., 2023), triose-phosphate isomerases(Romero-Romero et al., 2024), malate dehydrogenases(Johnson et al., 2025), or nucleases(Ivančić et al., 2025). Together, these studies illustrate the ability of pLM-based approaches to expand natural protein family repertoires with artificial sequences. However, pLMs have not yet been explored for small-molecule binder design, and most state-of-the-art pLMs are not trained as generators conditioned on specific functions. Crucially, when conditioning is supported, it has been typically limited broad labels (e.g., taxonomy tags(Madani et al., 2020) or EC/GO classes(Munsamy et al., 2024)) and training set instances. This setup does not allow generalisation to user-defined molecules or target definitions at the molecular level, which will be crucial to fully reach user-defined controllable protein design. Therefore, it is crucial to evaluate generators with respect to a specific target instance.

A widely successful strategy in NLP for such conditioning is to frame diverse tasks as text-to-text (sequence-to-sequence) problems, where an encoder transforms the input into a contextual representation and a decoder generates an output sequence conditioned on it. This paradigm has also inspired protein models that generate sequences conditioned on userdefined structures (Dauparas et al., 2022). However, protein design conditioned on a small molecule or substrate has not been explored to date. To address this gap, we provide a comprehensive perspective on the performance of brute-force protein language modeling for small-molecule binding design.

Here, we curate several ligand–protein datasets (with > 17*M* ligand-protein pairs) and develop several models (from 16*M* to 700*M* parameters) which take a target ligand as input and generate a binder-protein as output. We evaluate generalisation across data regimes, and benchmark architectures, model sizes, sampling schemes, input representations, and a pre-training strategy. We analyse generalisation across data regimes and observe a trade-off: settings with a low molecule/sequence ratio favour enhanced diversity during generation but reduce foldability, whereas the opposite yields a “retrieval-like behaviour” returning highly–foldable low–novelty sequences. Even for low–novelty generations, our results provide evidence of “ligand novelty” by discovering unseen protein–ligand interactions. These results highlight the potential and limitations of current datasets and training paradigms, which have an impact on the broader training of pLMs. To facilitate future work, we release our datasets and pretrained models on Hugging Face^1^ and our code on GitHub^2^.

## 2. Related Work

Recent generative-AI approaches to small-molecule binder design have been predominantly structure-centric, and many of the best-known demonstrations leverage diffusion models to generate ligand-compatible backbones. For example, RFdiffusion All-Atom (Krishna et al., 2024) generates backbones conditioned on a target structure and/or hotspot constraints (Fox et al., 2025), and has been applied to the design of digoxigenin, heme, and bilin binders. A common pipeline first generates a backbone containing a ligand-binding site, then derives an amino-acid sequence by inverse folding with methods such as ProteinMPNN, and finally ranks or refines candidates via re-folding or co-folding models such as AlphaFold2 (Baek et al., 2021), AlphaFold3 (Abramson et al., 2024), or Boltz (Passaro et al., 2025). In a complementary direction, frameworks such as BoltzDesign (Cho et al., 2025) directly optimize sequences (or designs) under a co-folding model by maximizing the likelihood of protein–ligand contacts under the model distribution. Beyond diffusion-based backbone generation, LigandMPNN (Dauparas et al., 2025) takes protein backbones together with docked molecules as input and designs sequences for the specified conformation, with experimental validation reported for the redesign of multiple ligand-binding proteins.

While these structure-first methods benefit from strong geometric inductive biases, their applicability depends on access to high-resolution protein–ligand complex structures (or reliable poses and binding-site definitions). This motivates sequence-based alternatives: the breadth of accumulated sequence data is on the order of six orders of magnitude larger than the set of structurally resolved complexes, making approaches that operate directly on 1D sequences, such as protein language models (pLMs), particularly appealing(Richardson et al., 2023).

Prior work has shown that pLMs can incorporate useful control signals during training. Decoder-only models such as ProGen condition generation on taxonomic and functional keyword tags, while ZymCTRL (Munsamy et al., 2024) conditions on Enzyme Commission (EC) labels to generate sequences matching a desired catalytic reaction class. These models can generate active enzymes, but their controllability is largely limited to the EC numbers and GO terms represented in their training sets, which restricts generalisation to unseen functions. Other efforts have targeted covalent binding prediction: T5ProtChem (Kelly et al., 2025) trains an encoder–decoder model that takes protein–ligand pairs as input and outputs a SMILES string and binding position, together with a probability of covalent binding and can also be used for protein function classification and reaction prediction.

Encoder–decoder architectures have therefore been widely explored for structure-conditioned sequence design (Ingraham et al., 2019; Dauparas et al., 2022), however, the application of sequence-to-sequence conditioning directly on small molecules for protein generation remains completely underexplored.

## 3. Methods

### 3.1. Dataset

We created two datasets for training representing opposing data-availability regimes. The Binder dataset contains more unique molecules than proteins, whereas the Substrate dataset maps multiple unique proteins to each molecule (Figure 1b, and Figure A.2)

**Figure 1.**
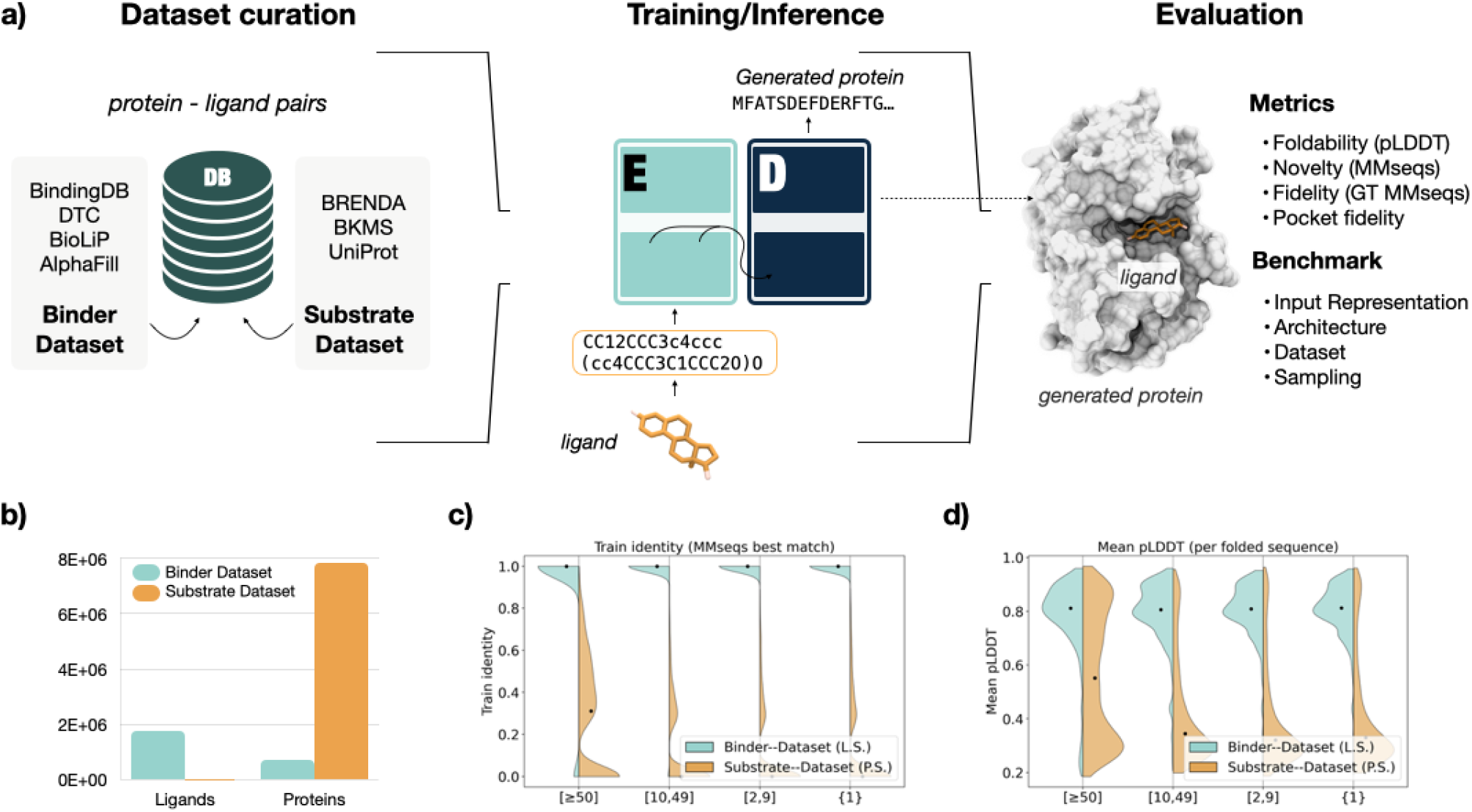
**(a)** Pipeline Summary **(b)** Dataset sizes **(c)** Protein novelty for the [≥50] train evaluation split for the Ligand–Sampling model trained on the Binder–Dataset and Pair–Sampling model trained on the Substrate–Dataset, representing the two extremes of the distribution **(d)** pLDDT evaluation for the same models and split.

#### Binder–Dataset

The training dataset was assembled by aggregating curated data from BindingDB (Liu et al., 2025), Drug Target Commons (DTC) (Tanoli et al., 2018), BioLiP (Yang et al., 2012), and AlphaFill (Hekkelman et al., 2023). Records missing identifiers, measurements, or displaying non-exact affinity relations were discarded. Ligand notation was standardised converting InChi codes to canonical SMILES. For DTC, compound/target identifiers were resolved via ChEMBL and UniProt to obtain ligand InChI and target sequences, respectively. Following common practice in literature, we retained only interactions with a reported affinity (IC50, Kd, or Ki) ≤10 *µ*M, except for BioLip which we include fully. Finally, duplicated protein–ligand pairs were removed, and the set of binding proteins for each ligand was filtered to 90% identity with MMSeqs2 (Steinegger & Söding, 2017). The final dataset contains ∼1.8M unique ligand SMILES, ∼774k unique protein sequences and ∼10M total ligand–protein pairs, averaging 5.55 proteins per ligand (and 2.0 median). The dataset follows a long-tailed distribution, with 0.25% most annotated summing up to 51% of the total pairs, corresponding to promiscuous ligands that interact with thousands of proteins each. This is addressed using sampling strategies.

#### Substrate–Dataset

As an additional experiment, we also retrieved data from enzyme datasets. Given a protein known to catalyse a reaction, we can assume the reactants also bind the protein. We extract reaction–enzyme pairs from Rhea (Bansal et al., 2022), BKMS (Lang et al., 2011), Uniprot (Bateman et al., 2025) and BRENDA (Chang et al., 2021). Then, from the substrate side of each one, we extract candidate ligands and filter by RDKit validity, heavy-atom count ∈ [5, 50], and remove a small list of trivial species and ubiquitous metabolites (e.g., acetate/succinate/ATP). We discard reactions containing unspecified (wildcard) atoms, and allow at most a 2 hydrogen / 1 oxygen mismatch between substrate and product atoms (to account for common redox / protonation conventions / loss of a water molecule). SMILES are canonicalised, allowing stereo-chemistry when available. The final training set contains 4015 unique ligands, ∼7.8M protein sequences, and a total of ∼17M pairs, averaging ∼3600 proteins per ligand. Additionally, we also define Extended–dataset as the union of Substrate–dataset and the Binder–dataset.

#### SAIR

We create a third dataset for testing, retrieved from the Structurally Augmented IC50 Repository (SAIR) (Lemos et al., 2025) and curated by discarding molecules failing basic chemistry and size constraints (valence, heavy-atom count, molecular weight, and rotatable-bond limits). We then extract per-residue heavy-atom coordinates for the polymer chain and all non-polymer components as candidate ligands/cofactors. Non-polymer components are clustered using contact-based merging: residue–ligand contacts are defined by an atom-pair distance threshold given by the sum of van der Waals radii plus a margin of 0.5Å (as computed in BioLip), then components are merged and considered in the same pocket when their contacting-residue sets have Jaccard overlap *≥* 0.30 and their minimum inter-component atom distance is ≤4Å . Like this, for each protein-ligand pair, we obtain a list of the residues in contact with the ligand and index them with their pocket identity, allowing to identify multiple pockets for the same ligand if available. The resulting dataset collects *∼*735K unique ligands, *∼*5k unique protein sequences and *∼*1M total ligand–protein pairs, averaging *∼*1.4 proteins per ligand.

### 3.2 Model and training

Models were trained with a cross-entropy loss between the input molecule *x*^′^ (text-based tokenized) and the target enzyme sequence *y* = (*y*_1_, *…, y*_*T*_ ) (amino acid tokens). Full details in Section A.1. We performed experiments with the architectures summarised in Table A.1. The main encoder– decoder followed the original T5 architecture (Raffel et al., 2020), using the T5-Base configuration, unless otherwise stated. Section 4.4.2 explores the integration of a LLama3 model (Grattafiori et al., 2024) as decoder, with pretrained weights from large scale protein language modelling. This pre-trained model was trained on 43M protein sequences obtained from Uniref90 and Gigaref (small molecule binders and non-binders). To condition the generation on an encoder representation, cross-attention layers were added to the decoder after each self-attention one. The input embedding matrix and the language modelling (LM) head were reshaped to the size of the new vocabulary. Then, training was performed in two stages. Stage–1 trained the encoder plus decoder’s cross-attention, input embeddings and LM head, aiming to achieve a local minima that reuses decoder representations. Then, Stage–2 trained all parameters.

#### Sampling

To compensate for the effect of distribution imbalance (Figure 1), we devise a *Unique–Ligand* sampling strategy, where each input (ligand) SMILES is sampled once per epoch and the target output protein is chosen at random (without replacement) from its binding list. This follows the same general principle used in previous large-scale pLM training to improve sample efficiency by preventing redundant sequence groups from dominating optimisation (Cheng et al., 2024; Meier et al., 2021). When unique-input sampling is performed, the number of epochs is increased to keep the number of training steps comparable to naive Pair sampling, given that each epoch only samples one solution per input. Comparative results between the two sampling strategies are provided in Table 1.

**Table 1.** Main model evaluation trained with Ligand and Pair sampling at equal number of steps.

| | SPLIT RANGE | MEAN±STD PLDDT | GT MMSEQS (ACC. \| MEAN) |
| --- | --- | --- | --- |
| **LIGAND SAMPLING** |  |  |  |
| **TRAIN** | [50, 24570] | 77.67±15.14 | 78.37% \| 0.30±0.11 |
| | [10, 49] | 79.05±12.23 | 92.46% \| 0.77±0.16 |
| | [2, 9] | 79.70±11.27 | 72.51% \| 0.57±0.10 |
| | {1} | 79.91±10.49 | 46.34% \| 0.40±0.06 |
| | SAIR | 79.07±10.68 | 38.31% \| 0.32±0.04 |
| | SAIR POCKET | – | 39.51% \| 0.33±0.05 |
| **TEST** | [50, 346253] | 75.94±18.35 | 46.17% \| 0.23±0.07 |
| | [10, 49] | 79.12±12.78 | 90.89% \| 0.71±0.19 |
| | [2, 9] | 79.75±12.05 | 51.29% \| 0.39±0.08 |
| | {1} | 78.41±13.14 | 32.86% \| 0.28±0.04 |
| | SAIR | 79.36±10.42 | 38.82% \| 0.33±0.05 |
| | SAIR POCKET | – | 39.56% \| 0.32±0.05 |
| **PAIR SAMPLING** |  |  |  |
| **TRAIN** | [50, 24570] | 65.66±24.33 | 72.57% \| 0.30±0.10 |
| | [10, 49] | 77.79±14.31 | 93.50% \| 0.79±0.17 |
| | [2, 9] | 77.24±15.25 | 45.77% \| 0.41±0.04 |
| | {1} | 77.14±15.17 | 17.50% \| 0.15±0.01 |
| **TEST** | [50, 346253] | 55.11±26.19 | 37.39% \| 0.19±0.06 |
| | [10, 49] | 76.49±16.37 | 89.94% \| 0.77±0.15 |
| | [2, 9] | 76.59±16.34 | 34.53% \| 0.31±0.03 |
| | {1} | 73.76±19.25 | 9.49% \| 0.08±0.01 |

#### Tokenizer

In our database, we represent ligands as SMILES (Weininger, 1988) strings (or SELFIES (Krenn et al., 2020) for the input comparison experiments), while output proteins are represented using the standard 20–aminoacid alphabet. We developed a tokenizer tailored for ligand-conditioned protein generation, which we release to the community. Full details available in Section A.2.

### 3.3 Metrics

Protein generation (25 generations per input ligand; top-k sampling with k=15 and temperature t=1) was evaluated both for the test and training sets. Test set ligands are not present in the training set, while associated proteins can overlap. The two sets were further divided grouping by number of annotated proteins per ligand, in ranges: [50,∞), [10, 49], [2, 9], {1} . This stratification enables evaluation across regimes of annotation density and label ambiguity (from highly multimodal to near-singleton supervision). We selected 200 unique ligands per split.

To evaluate structural reliability of the model’s designs, we predicted their folded structure using ESMFold (Lin et al., 2021). This approach has been common practice in the protein design community, given the correlation between pLDDT and the likelihood of the sequence being ordered (Tunyasuvunakool et al., 2021). To quantify the similarity of predicted proteins to the training set, for each prediction, we compute alignment–based similarity through MMSeqs2 against all training samples (Train Id.). This quantifies the novelty of a sequence, indicating to what extent the protein is retrieved from the training distribution.

Finally, we assess generalisation leveraging the ground-truth (GT) annotation. Given that test ligands are not seen during training, a prediction that matches an annotated GT binder (or a close homolog) provides evidence that the model generated a plausible binder for an unseen ligand. On the contrary, not matching GT is no guarantee of an invalid solution. For GT matches, generalisation is arguably proportional to how different the test ligand is to the training examples. Using MMSeqs2 search, non-exact matches will also be identified, as sequences with few amino acid mutations have a high likelihood of converging to a similar fold. We compute GT accuracy (GT Acc.) by selecting the maximum identity match among all generated proteins per input (25), and averaging across all test samples. We also report the mean and standard deviation of all matches.

## 4. Results

Results in Table 1, under “Ligand Sampling”, present the performance of our best scoring model, which is trained on the Binder–dataset with Unique–Ligand sampling. Mean pLDDT indicates that the network yields foldable sequences across different splits, decaying only slightly for the most promiscuous ligands. GT accuracy demonstrates non-trivial success in the retrieval of annotated binders across most ligand strata; particularly in the moderately annotated regime (90.89% GT acc.). In contrast, GT retrieval decreases for the most promiscuous test ligands (46.17% GT acc.), which likely reflects a harder multi-modal distribution following a less deterministic pattern. For ligands that only map to a single protein in the dataset, performance is lower but non-negligible (32.86% GT acc.).

Leveraging the pocket annotations provided by the SAIR split (Section 3), we extract the ground-truth (GT) pocket residues and compute sequence identity specifically over the binding site. Then we compare it to the identity towards the full (GT) protein. We sample 400 unique ligands for each SAIR evaluation set, one with ligands seen during training and another with hold-outs (protein/molecule ratio presented in Figure A.2). Results in Table 1 indicate a closely similar score for pocket (“SAIR Pocket”) vs full protein matches (“SAIR”), with “Pocket” being slightly higher. Binding specificity is primarily determined by residues in the binding pocket; therefore, these results suggest that GT matches are not driven solely by non-functional sequence regions but are also supported by similarity at the binding site.

Finally, as previously explained, Unique–Ligand sampling is used to prevent the over-sampling of heavily annotated ligands. Table 1 presents results for the same model under regular Pair–Sampling, using the same number of training steps. Ligand sampling improves both structural confidence and GT retrieval relative to pair sampling, with the largest gains appearing in sparsely supervised regimes (e.g., test singletons and the [2, 9] stratum).

### 4.1. Ligand novelty

To assess generalisation, we measure the novelty of test ligands relative to the training set. Specifically, we quantify how dissimilar a target ligand is from ligands associated with the closest training proteins in sequence space. Correct generations that match the GT while involving ligands far from any training example indicate generalisation beyond memorisation of observed protein–ligand pairs.

For each generated protein, we compute its MMseqs identity versus all train samples and select its nearest neighbours (all maximally similar samples, or the top 50 if fewer). Instead of regular identity *id*, given alignment length *a* and target lenght *t* we define an effective identity 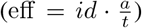, to favour matches of larger coverage. We retain the set of closest training proteins within a tolerance: eff ≥eff_max_ −0.05 (with the exception of GT sequences that are always included if they are listed as a match). We then pool all ligands paired with these proteins in the training set and measure their similarity towards the test ligand by means of Tanimoto similarity with Extended Connectivity Fingerprints (Rogers & Hahn, 2010) (radius 2, 2048 bits). Finally, the maximum Tanimoto similarity (MTS) is reported. This value indicates the following: If the generated protein closely matches a training example *E*, the reported value is the maximum chemical similarity found for a ligand associated to *E* and the test ligand.

Figure 2.A plots GT identity versus MTS for each generated sample with the main model on the [≥50] test split (results for the rest of splits available in supplementary Section B). We observe that MTS spans a broad range even when the generated protein closely matches an annotated GT binder. Successful GT retrievals (high GT identity) occur not only for ligands that are near-duplicates of ligands seen with similar training proteins, but also for ligands with low chemical similarity to any ligand in the neighbourhood. We interpret these cases as ligand-level generalisation: the model recovers an annotated protein (or close homolog) for a ligand that is chemically dissimilar to ligands observed in the training neighbourhood of that protein family.

**Figure 2.**
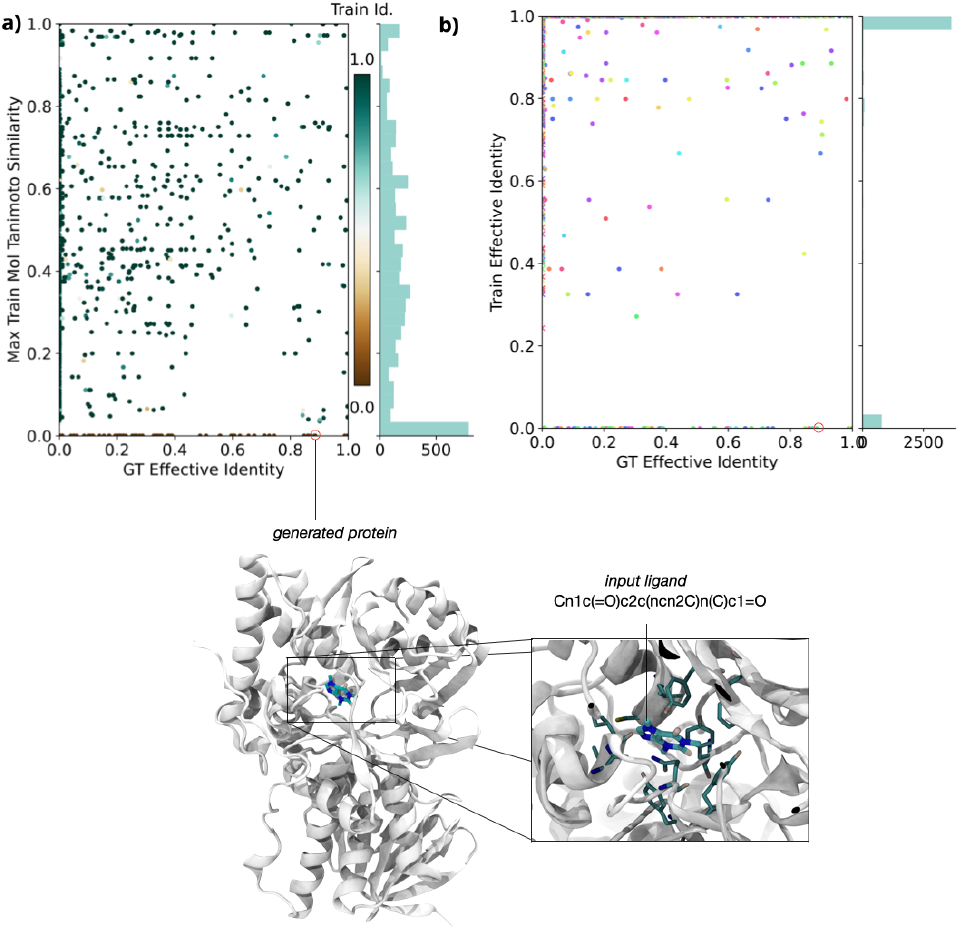
**(a)** MTS versus GT eff id. per generation in the [≥50] test split. **(b)** Train eff id. versus GT eff id. on the same split. Bottom: Co-folding by Boltz2 of a high GT identity generation with no training match (0.97 complex pLDDT, 0.85 iPTM).

To contextualise how much of the GT retrieval can be explained by chemical-similarity retrieval, Section B.2 computes evaluates GT matching with a Tanimoto retrieval baseline.

### 4.2. Protein novelty

We compute MMSeqs2 similarity of generated sequences against the training set (Train Id.). For the model trained with Pair Sampling, training hits go from ∼52–95% depending on stratum, while Ligand sampling increases to ∼85–99% (full results in Supplementary Figure A.1). The distribution can be visualised in Figure 1.C. This same figure compares the distribution to the one obtained with the Substrate–Dataset.

Figure 2.B visualizes the relationship between GT retrieval and protein-level novelty by plotting the effective identity of each generation to the GT (x-axis) and to the closest training neighbour (y-axis). Cases where a generation matches the GT more closely than any training sequence provide a direct signature of protein-level generalisation: the model recovers a binder sequence that is closer to the held-out annotation than to the training distribution. Such events are expected to be rare, since they require the GT binder itself to be sufficiently distant from training proteins, which occurs infrequently in natural datasets. Nevertheless, the presence of such examples indicates that the model can move beyond nearest-neighbour retrieval and generate sequences aligned with held-out binders. Figure 2 displays the co-folding prediction of a generation with 0.88 GT effective identity (0.99 target coverage) and no training match, which was targeting the caffeine molecule. The co-folding was computed with Boltz2 using MSA templates, which predicts successful binding with high confidence (0.97 pLDDT and 0.85 iPTM). Therefore, despite the absence of caffeine binders in training, the model generated a sequence with high likelihood of binding according to Boltz2, while remaining 10% divergent from the closest GT annotation.

As a final analysis, we study the influence of generation temperature in sequence novelty. The model learned to reproduce low entropy distributions; therefore, we test whether higher sampling temperatures can increase such entropy and increase novelty. Table 2 does not support the hypothesis: increasing temperature and *k* in *top-k* sampling reduces the fraction of generations that match any training sequence, but the identity of the remaining matches remains similar. Together with the reduction in GT matching, this suggests off-target generations rather than useful novelty. Following the same trend, lowering temperature to *<* 1 values increases GT matching. This is coherent with a model that learnt a narrow output distribution and rapidly falls out of distribution when deviating from the greedy decoding.

**Table 2.** Effect of sampling temperature (*T* ) and top-*k* (*K*) on SAIR evaluation.

| $T$ | $K$ | TRAIN ID. | MATCHES | GT ACC. | MEAN |
| --- | --- | --- | --- | --- | --- |
| 0.8 | 15 | $1.00 \pm 0.01$ | 99.1% | 40.48% | $0.34 \pm 0.04$ |
| 1.0 | 15 | $1.00 \pm 0.01$ | 98.2% | 38.82% | $0.33 \pm 0.05$ |
| 1.4 | 50 | $1.00 \pm 0.04$ | 91.1% | 35.23% | $0.30 \pm 0.04$ |
| 1.6 | 50 | $0.99 \pm 0.06$ | 81.0% | 31.40% | $0.27 \pm 0.04$ |
| 1.6 | 100 | $0.98 \pm 0.06$ | 78.4% | 31.19% | $0.27 \pm 0.03$ |
| 1.8 | 50 | $0.97 \pm 0.08$ | 68.4% | 26.86% | $0.24 \pm 0.03$ |
| 2.0 | 50 | $0.93 \pm 0.11$ | 51.3% | 18.86% | $0.17 \pm 0.02$ |

### 4.3. Substrate–Dataset results

Using the Substrate–Dataset (defined in Section 3) we train an identical model on this alternative data distribution. Figure 1.C and 1.D compare the Train Id. and pLDDT distributions (respectively) of this model to the one trained on the Binder–Dataset, revealing substantial differences between the two. As depicted in Figure 1.B, the Substrate–Dataset contains orders of magnitude fewer ligands, and orders of magnitude more proteins, providing a wide distribution of outputs for each input. This causes the model to learn a distribution of higher entropy per input, closer to what it is observed in non-conditional pLMs when learning the full distribution of natural proteins.

Notice that the higher variability in generation comes at the cost of lower pLDDT values, which is expected, as retrieving known proteins from training set is arguably an upper bound for foldability, while deviating from the natural distribution is known to reduce it. Table 3 reports the complete results, including Ligand and Pair sampling. As expected, when Ligand Sampling allows the lower entropy regions ([2, 9], {1}) to be trained on as much as the largest ones, they move closer to a retrieval behaviour.

**Table 3.**
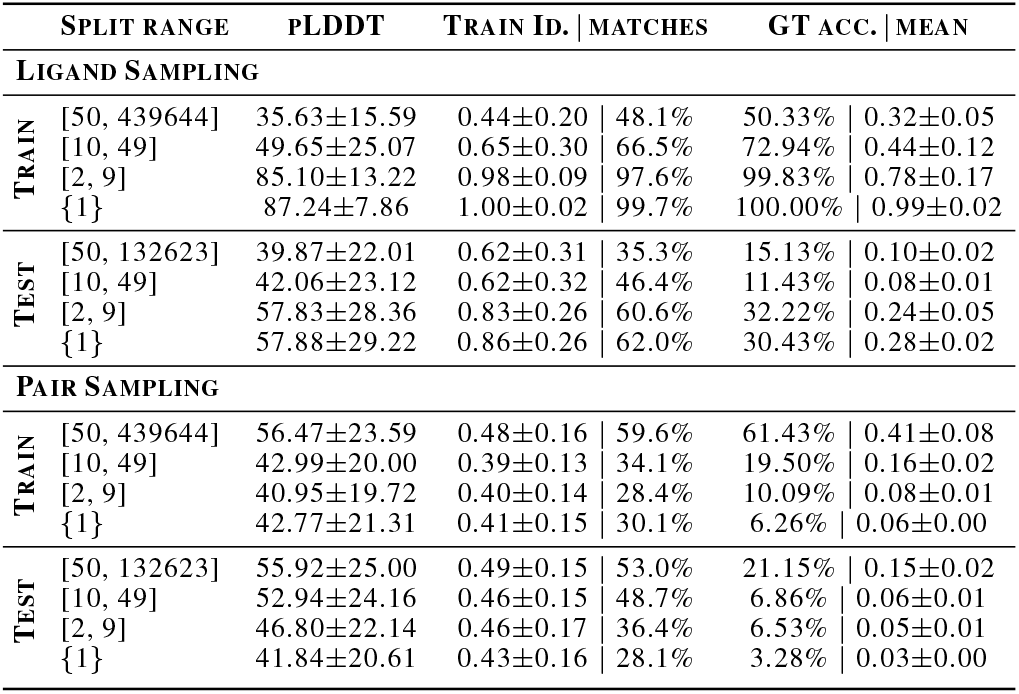
Evaluation on the substrate–enzyme dataset.

|  | SPLIT RANGE | PLDDT | TRAIN ID. | MATCHES | GT ACC. | MEAN |
| --- | --- | --- | --- | --- | --- | --- |
| **LIGAND SAMPLING** |  |  |  |  |  |  |
| TRAIN | [50, 439644] | 35.63±15.59 | 0.44±0.20 | 48.1% | 50.33% | 0.32±0.05 |
|  | [10, 49] | 49.65±25.07 | 0.65±0.30 | 66.5% | 72.94% | 0.44±0.12 |
|  | [2, 9] | 85.10±13.22 | 0.98±0.09 | 97.6% | 99.83% | 0.78±0.17 |
|  | {1} | 87.24±7.86 | 1.00±0.02 | 99.7% | 100.00% | 0.99±0.02 |
| TEST | [50, 132623] | 39.87±22.01 | 0.62±0.31 | 35.3% | 15.13% | 0.10±0.02 |
|  | [10, 49] | 42.06±23.12 | 0.62±0.32 | 46.4% | 11.43% | 0.08±0.01 |
|  | [2, 9] | 57.83±28.36 | 0.83±0.26 | 60.6% | 32.22% | 0.24±0.05 |
|  | {1} | 57.88±29.22 | 0.86±0.26 | 62.0% | 30.43% | 0.28±0.02 |
| **PAIR SAMPLING** |  |  |  |  |  |  |
| TRAIN | [50, 439644] | 56.47±23.59 | 0.48±0.16 | 59.6% | 61.43% | 0.41±0.08 |
|  | [10, 49] | 42.99±20.00 | 0.39±0.13 | 34.1% | 19.50% | 0.16±0.02 |
|  | [2, 9] | 40.95±19.72 | 0.40±0.14 | 28.4% | 10.09% | 0.08±0.01 |
|  | {1} | 42.77±21.31 | 0.41±0.15 | 30.1% | 6.26% | 0.06±0.00 |
| TEST | [50, 132623] | 55.92±25.00 | 0.49±0.15 | 53.0% | 21.15% | 0.15±0.02 |
|  | [10, 49] | 52.94±24.16 | 0.46±0.15 | 48.7% | 6.86% | 0.06±0.01 |
|  | [2, 9] | 46.80±22.14 | 0.46±0.17 | 36.4% | 6.53% | 0.05±0.01 |
|  | {1} | 41.84±20.61 | 0.43±0.16 | 28.1% | 3.28% | 0.03±0.00 |

The combination of Substrate and Binder datasets yielded similar results to training exclusively on the Binder–Dataset, without showing improvement. Results are available in Supplementary C.

### 4.4. Architecture experiments

#### 4.4.1. Decoder-only architecture

Following the procedure described in Section 3, we trained a decoder-only model to explore the impact of the architectural choice in performance. Results in Table 4 show a lower test GT accuracy along with decreased average pLDDT compared to the encoder-decoder version.

**Table 4.** Encoder–decoder (t5-tiny) versus decoder-only (GPT2) in a common 100-sample test set. Both models were trained for 2 epochs with pair sampling.

| MODEL | PARAMS (M) | PLDDT | GT ACC. | MEAN |
| --- | --- | --- | --- | --- |
| ENCODER–DECODER | 7.6 | 65.20±25.18 | 22.24% | 0.21±0.01 |
| DECODER–ONLY | 14.6 | 61.67±11.37 | 17.01% | 0.16±0.02 |

It is important to note that these results are obtained for a specific dataset and architecture size with limited hyperparameter tuning. Nevertheless, given the proven data and computing efficiency of encoder-decoder architectures for conditioning tasks (Tay et al., 2022; Elfeki et al., 2025), the results confirm that they are easier to train for our task.

#### 4.4.2. Protein language modelling pretraining

The performance of the pretrained Llama3 model was evaluated in the SAIR test set and compared to the baseline t5 one, both trained in the main Binder–dataset and also in the Extended–dataset. Baseline models were trained from scratch for 45k steps. Pretrained ones employed 10k in Stage–1 (frozen decoder, as defined in Section 3) and 45k steps in Stage–2, where all parameters are trained. All models used a common batch size (1152), consequently, they were exposed to the same number of examples during unconstrained training, avoiding an unfair advantage for the baseline ones.

Results show how the model trained from scratch outperforms the pretrained one, both in pLDDT and GT matching. Regarding the train identity metric, it seems to indicate that the pretrained model did not generate a richer output distribution. Same as with the baseline model, generations finding alignments have maximum identity. This indicates that our training pipeline did not manage to preserve the richer distribution in the decoder, which converged to the same solution as the baseline. Additionally, lower performance indicates either that pretrained initialisation hampered convergence or that the hyper-parameters, which we kept the same as for the baseline, were suboptimal for this approach. It remains an open question whether additional techniques can be used to preserve the original distribution of the decoder while achieving competitive results.

#### 4.4.3. Parameter scaling

Performance across model sizes was computed for the tiny, base, and large configurations of the T5 family. Table 5 presents average pLDDT for the SAIR test set. Consistent with literature, we observe diminishing performance returns, hinting to logarithmic improvement over the number of parameters. In particular, the ∼700M configuration is only slightly superior to the ∼200M one (80.14 vs 79.36 average pLDDT). Hence, we select the ∼200M for the experiments in this work.

**Table 5.** Effect of model size on SAIR test pLDDT. t5-Tiny and T5-base were trained for 45K steps while T5-Large for 40K.

| MODEL | PARAMS (M) | PLDDT |
| --- | --- | --- |
| T5–TINY | 7.62 | 47.00±25.56 |
| T5–BASE | 199.05 | 79.36±10.42 |
| T5–LARGE | 705.86 | 80.14±9.65 |

**Table 6.** Pretrained-llama models vs baseline in SAIR test set. “P.” stands for pretrained, Binder for model trained on the Binder dataset and Extended for the joint Binder + Substrate dataset.

| MODEL | PLDDT | TRAIN ID. | MATCHES | GT ACC. | MEAN |
| --- | --- | --- | --- | --- | --- |
| BINDER | 79.07±10.68 | 1.00±0.01 | 99.8% | 38.82% | 0.33±0.05 |
| EXTENDED | 70.62±22.78 | 0.99±0.02 | 66.6% | 31.24% | 0.27±0.03 |
| P. BINDER | 74.13±15.42 | 1.00±0.01 | 76.3% | 19.85% | 0.18±0.02 |
| P. EXTENDED | 57.22±16.70 | 1.00±0.02 | 71.0% | 27.30% | 0.25±0.02 |

### 4.5. Input format experiments

To assess the relevance of input formatting, here we report results with SELFIES (Self-Referencing Embedded Strings) (Krenn et al., 2020), an alternative molecular string representation. Unlike SMILES, in SELFIES every syntactically valid sequence corresponds to a valid chemical structure, aiming to simplify the space of solutions for generative models. This comes at the cost of creating longer strings. Table 7 presents results of identical models trained with SMILES vs SELFIES. We do not find benefit in this alternative. We hypothesise that the advantage of an “always-valid” space is relevant when used in the generation output, but potentially unnecessary at the input. We refer readers to studies comparing the two representations (Skinnider, 2024; Flam-Shepherd et al., 2022; Leon et al., 2024).

**Table 7.**
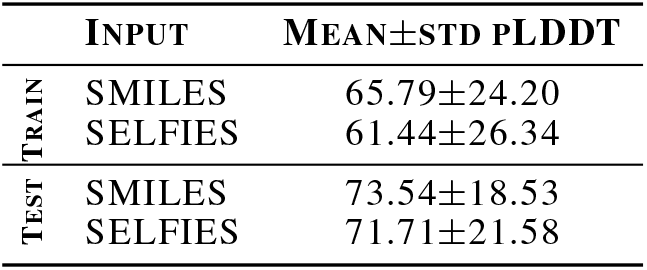
pLDDT results for identical t5-tiny models using SMILES or SELFIES as ligand representation. All models were trained for 5 epochs and evaluated in the same sets of 100 examples each.

### 4.6. Maximal distant test set

As an additional test of the generalisation capabilities of the model, we construct a test set that is maximally distant from training in protein sequence space, while associated ligands are also held out from training. Full algorithmic details available in Section D.

Table 8 presents results of the evaluation in this set, showing a sharp drop in GT identity. Strata with the lower annotation volume retain some identity to GT and high pLDDT, while splits ∈ [≥10] collapse, yielding sequences of lower fold-ability. Train identity remains high; therefore, the model is still retrieving seen protein families rather than adapting to the held-out distant ones. This behaviour is expected under our construction: when ground-truth binders are forced to be far in sequence space, any training-like output will score poorly against GT by sequence similarity, even if it were a plausible binder for the ligand. This test confirms that retrieval behaviour persists when prompted out of distribution. The validity of output sequences has a much lower likelihood of correctness in this regime; still, these results are not by themselves conclusive evidence of non-binding or overfitting, since alternative binder families may exist.

**Table 8.**
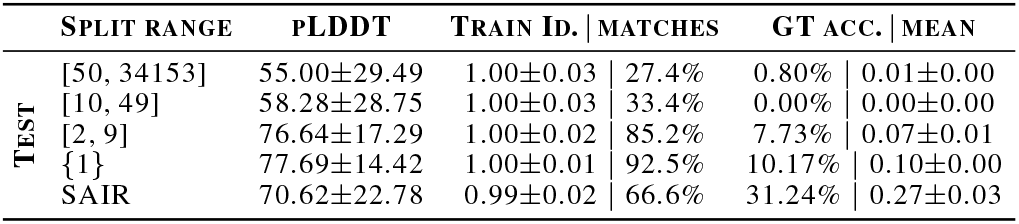
Maximally distant test set on Extended–Dataset. SAIR split remains the same as used for previous experiments.

## 5. Discussion and conclusions

We benchmarked ligand-conditioned sequence-to-sequence pLMs for small-molecule binder design using purely textual inputs, with the goal of understanding when brute-force conditional language modeling can propose plausible binders. We find that available annotation for most ligands is sparse, pushing the model towards a retrieval-like behaviour. This behaviour yields low novelty at the protein level, but as supported by the results on held-out molecules, it has potential for the discovery of new protein-ligand interactions.

When a ligand is paired with a narrow set of sequences, the cross-entropy optimum concentrates probability mass on a small number of modes; this makes it possible to minimise the loss by memorising and retrieving previously seen proteins or close homologs. When supervision for a ligand spans a broader set of binders (high conditional entropy), the model is forced to represent a richer conditional distribution, moving closer towards the behaviour of unconditional pLMs, which represent the extreme of this trade-off by sharing the same input for all sequences. Therefore, higher generalisation will be expected for ligands with rich annotation. Among them, as suggested by the lower performance of the [≥50] strata, GT will be most informative when the ligand is not excessively promiscuous, else, the output distribution will be highly multi–modal, making convergence harder, and probably requiring an even larger volume of annotation. This lesson helps to anticipate results based on data distribution, and hints at the current ceiling for language modelling on binder datasets, which is limited by the sparsity of annotation.

Ligand novelty analysis demonstrated how the model can generate a protein matching the GT (or close homolog) without seeing examples of such protein family being associated to chemically similar ligands. This requires a general understanding of chemical compatibility and binding activity, which allows to “fill the gaps” and understand which proteins could bind a newly found ligand. As discussed, these results are most often obtained by reproducing sequences seen during training, still we argue there is value in finding novel uses for known proteins. Additionally, we also find few cases where generation diverged from training but still proposed a plausible binder, such as the caffeine binder, validated by Boltz2 co-folding. Therefore, the model can be used to yield multiple binder proposals (in a matter of seconds) and then select the most promising ones for experimental validation. Filtering tests can include, among others, co-folding, docking or search against protein databases. We consider this post-processing to be an important component of future pipelines, especially since GT-based metrics are an imperfect proxy for binding success. It is of interest for future work to characterise success with higher fidelity measures and experimental validation.

To the best of our knowledge, pLM conditioning had not been yet studied for targets outside the training set, hence we believe an initial baseline was necessary. Here, we benchmark architectures, model sizes, generation parameters, input formats, and pLM pretraining. Together with the curated datasets, and evaluation pipelines, we hope to provide the necessary tools to lower the barrier for the exploration of ligand-conditioned protein generation.

Looking forward, the target for general–purpose AI binder design should be to push novelty, designing de-novo proteins which diverge from known sequences, while demonstrating high success rate in lab trials. As discussed, the role of pLMs in this development is tied to databases. Scaling text-based datasets is a way forward, but remains expensive. Alternatives include using richer annotation, such as binding affinity, or resorting to multi-modality with structural data. In addition, inductive biases, such as physics–based computations, might also prove necessary.

## Acknowledgements

We thank the members of the Ferruz Lab at CRG for their most valuable feedback and insights, with a special mention to Filippo Stocco for his assistance with co-folding. NF acknowledges support from the Ramón y Cajal Fellowship(RYC2021-034367-I) funded by MI-CIU/AEI/10.13039/501100011033 and the European Union NextGeneration EU/PRTR, and from the European Union’s Horizon Europe program under grant agreement No 101165231. N.F and A.V acknowledge support by the European Research Council (ERC) grant ATHENA (ERC-ST-2024, Grant agreement 101165231). J.C acknowledges support from MINECO Transmisiones project PLEC2023-010243. We acknowledge support of the Spanish Ministry of Science and Innovation through the Centro de Excelencia Severo Ochoa (CEX2020-001049-S, MCIN/AEI /10.13039/501100011033), and the Generalitat de Catalunya through the CERCA programme. For computing resources, authors gratefully acknowledge the scientific support and HPC resources provided by the Erlangen National High-Performance Computing Center (NHR@FAU) of the Friedrich-Alexander-Universität Erlangen-Nürnberg (FAU) under the NHR project b114cb (UID 210235). Federal and Bavarian state authorities provide NHR funding. NHR@FAU hardware is partially funded by the German Research Foundation (DFG) - 440719683. Finally, we acknowledge EuroHPC for the JU Development Access grants EHPC-DEV-2025D03-066 and EHPC-DEV-2025D11-153 which granted access to the MareNostrum5 supercomputer hosted by Barcelona Supercomputing Center. Views and opinions expressed are, however, those of the author(s) only and do not necessarily reflect those of the European Union. Neither the European Union nor the granting authority can be held responsible for them.

## Impact Statement

The primary impact of this work is to provide rigorous foundation for instance-level conditioning in protein language models. Outcomes and insights achieved in this work assist researchers in building next-generation conditional pLMs and enable practitioners to triage candidate binders via downstream filtering (e.g., co-folding/docking) before any experimental work. There are no direct societal impacts from this work alone, as we do not report experimentally validated binders or deployable design protocols. In the long run, contributions to general purpose AI binder design aim to enable faster discovery for therapeutics, sustainability, or industrial purposes. The consequences of such developments are shared across all disciplines contributing to these ends.

## Appendix

### A. Model details

#### A.1. Training

Given a network parametrised by weights *θ*, the loss is defined as the cross-entropy between the input molecule *x*^′^ (text-based tokenized) and the target enzyme sequence *y* = (*y*_1_, *…, y*_*T*_ ) (amino acid tokens):

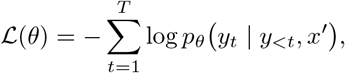

where *p*_*θ*_(*y*_*t*_ | *y*_<*t*_, *x*^′^) is the probability assigned by the model to token *y*_*t*_ given previous target tokens *y*_<*t*_ and the encoder output.

The main model was trained for 30 epochs with unique-input sampling. Hyperparameters consisted of: 2 × 10*e*^−4^ learning rate, cosine decay to 2 ×10*e*^−5^ (through 5 epochs), 3000 warm-up steps (3.3% of the total), 1152 batch size, and bf16 mixed precision. Alternative models reused these hyper-parameters unless otherwise stated. The training hardware consisted of 8 H100 Nvidia GPUs with 64Gb VRAM.

#### A.2. Tokenizer details

Our datasets represent ligands as SMILES strings while proteins are represented using the standard 20–amino-acid alphabet. To avoid ambiguity between chemical symbols and amino-acid letters (e.g., “C” denoting both carbon and cysteine), we define two separate vocabularies and use a dedicated tokenizer for each modality. Concretely, if the ligand tokenizer defines *n* tokens, we offset the indices of the protein tokenizer by *n*, ensuring that the two token spaces do not overlap.

The amino acid tokenizer has a token for each possible amino acid. In contrast, the SMILES tokenizer uses a “word-piece” tokenization algorithm, as the space of input characters and “chemical words” is much larger.

When tokenizing chemical notation, one can simply define a token per unique character in the dataset. Still, LLM literature has demonstrated how grouping tokens that frequently appear together improves computational efficiency and generalisation. That is because a single token prediction will yield a larger amount of information and because the model is allowed to have a dedicated embedding for combinations of characters, starting from a larger library of unique concepts. As described in previous work (Leon et al., 2024), applying tokenization algorithms such as BPE directly to SMILES, can generate invalid tokens, as atom notations such as “He” cannot be split as “H” + “e”. Therefore, as done in (Leon et al., 2024) with their APE tokenizer, we define a 2 stage algorithm, which first defines indivisible atom tokens, and then performs the BPE algorithm to join those that frequently co-occur. We name our algorithm ABPE, as it fully reproduces the original BPE algorithm for the second stage, unlike the APE algorithm.

#### A.3. Architectures

Table A.1 lists the architectural details of all models used in this work, differentiating encoder and decoder parameters, and specifying the size of the language modelling (LM) head, which depends on the vocabulary size.

**Table A.1.** Model architecture summary. For encoder–decoder models, we report encoder and decoder hyperparameters; when a value is identical for encoder and decoder we list it once, otherwise we use encoder |decoder (e| d). *P*_enc_ and *P*_dec_ denote the number of parameters (in millions) in the encoder/decoder, *P*_LM_ in the output head for a vocabulary size of 1069, and *P*_tot_ for the total.

| MODEL | $P_{\text{enc}}$ (M) | $P_{\text{dec}}$ (M) | $P_{\text{LM}}$ (M) | $P_{\text{tot}}$ (M) | $L$ | $H$ | $d_{\text{model}}$ | $d_{\text{ff}}$ | $d_{\text{kv}}$ |
| --- | --- | --- | --- | --- | --- | --- | --- | --- | --- |
| T5-TINY | 3.1 | 4.2 | 0.27 | 7.62 | 4 | 4 | 256 | 1024 | 64 |
| T5-BASE | 85.0 | 113.3 | 0.82 | 199.05 | 12 | 12 | 768 | 3072 | 64 |
| T5-LARGE | 302.0 | 402.7 | 1.09 | 705.86 | 24 | 16 | 1024 | 4096 | 64 |
| GPT2 (DECODER-ONLY) | — | 14.2 | 0.41 | 14.61 | 8 | 6 | 384 | 1536 | 64 |
| T5-BASE ENC + LLAMA DEC | 85.0 | 146.8 | 1.09 | 232.87 | 12\|10 | 12\|16 | 768\|1024 | 3072\|2048 | 64 |

**Figure A.1.**
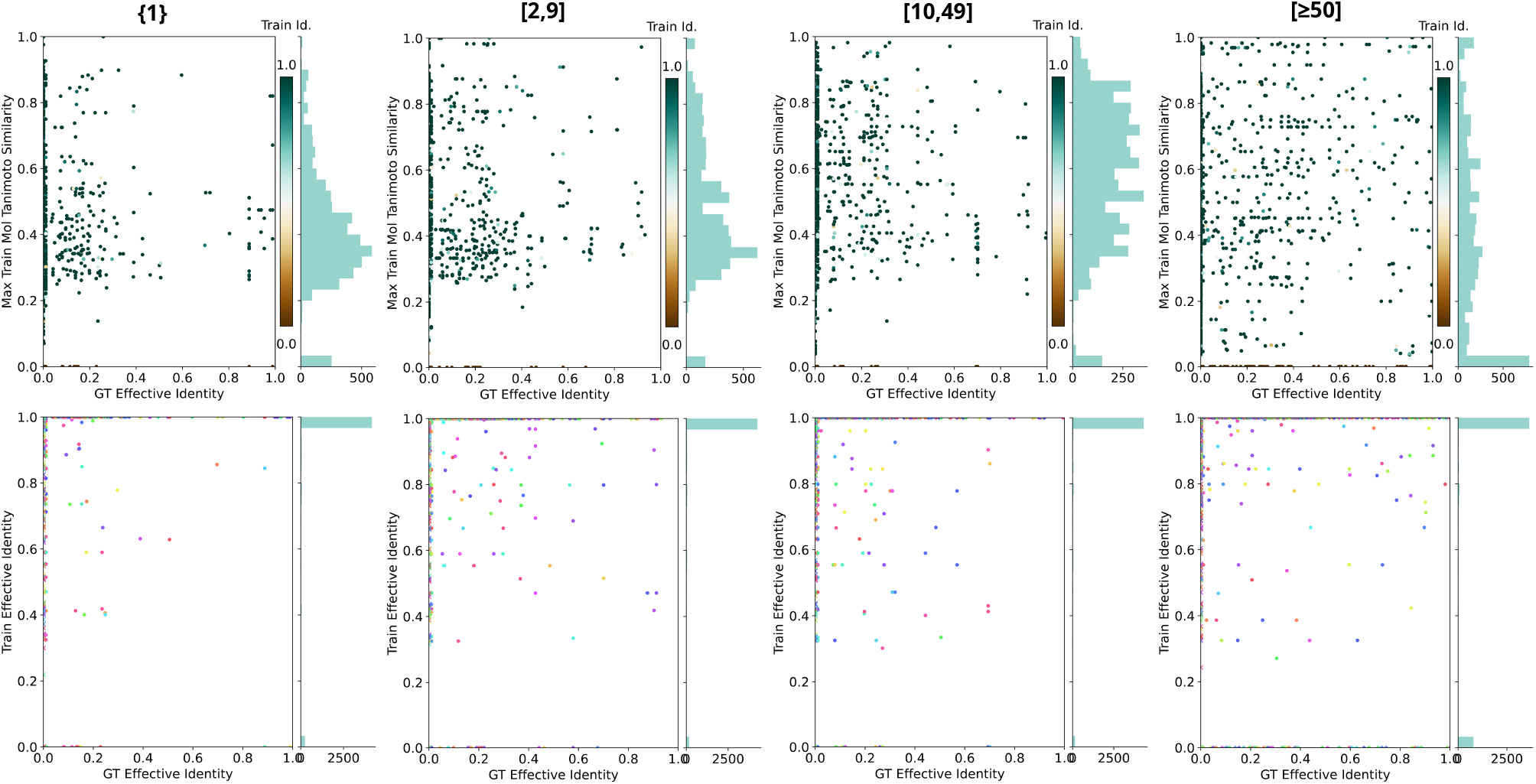
Ligand novelty graphs per test split.

### B. Complete novelty results

Figure A.1 provides the full results for GT identity vs MTS and Train identity. Notice how the MTS distribution becomes most widespread (and furthest from training) for the [≥50] split. This is consistent with intuition, as the most promiscuous ligands are most likely to be associated to widely different proteins, giving the model the opportunity to bind them with protein families which are not specialised in this specific chemical profile.

#### B.1. Protein Novelty

Regarding protein novelty, Figure A.1 Table A.2 presents the full results with Train Id. for each split.

#### B.2. Fingerprint Nearest Neighbour retrieval baseline

As a diagnostic baseline to probe dataset overlap and the sensitivity of the GT identity metric to retrieval, we implement fingerprint *k*-nearest-neighbour (FP-NN) retrieval. For each test ligand, we compute a Morgan fingerprint (ECFP4; radius 2, 2048 bits) and retrieve the most similar ligand from the training set by Tanimoto similarity. We then output the protein sequence associated with each retrieved training ligand. The resulting FASTA outputs are evaluated with the same GT MMSeqs pipeline used for model generations. This baseline is expected to recover an annotated binder family when chemically similar ligands exist in training and their associated proteins are the same or close homologs to the GT.

Table A.3 shows how FP-NN retrieves proteins matching the GT for a large fraction of test instances. Then, we use these results to quantify how often the model goes beyond simple retrieval. We compared to the model’s greedy decoding, to make a fair 1-shot comparison, and computed the percentage of targets for which the model generation was closer to the GT than the FP-NN protein. We also verify that in these cases, the model proposal differs substantially in GT matching identity from the retrieval one (Table A.3 Mean Δ_model>_ column). This reinforces the conclusions of the generalisation studies, which showed examples beyond retrieval.

In terms of average matching, FP-NN matches the GT protein more frequently than the model. This suggests that the network does an imperfect job of memorising all training proteins and matching them to suitable targets. It was seen how, in cases where retrieval is not enough, the model can go beyond, but the occurrence of these cases in the Binder-dataset is not high enough to make the global average higher than pure retrieval. The smallest margin is found in the [10, 49] stratum, the greedy decoding matches GT at 38.83% and FP-NN at 46.62%.

**Table A.2.**
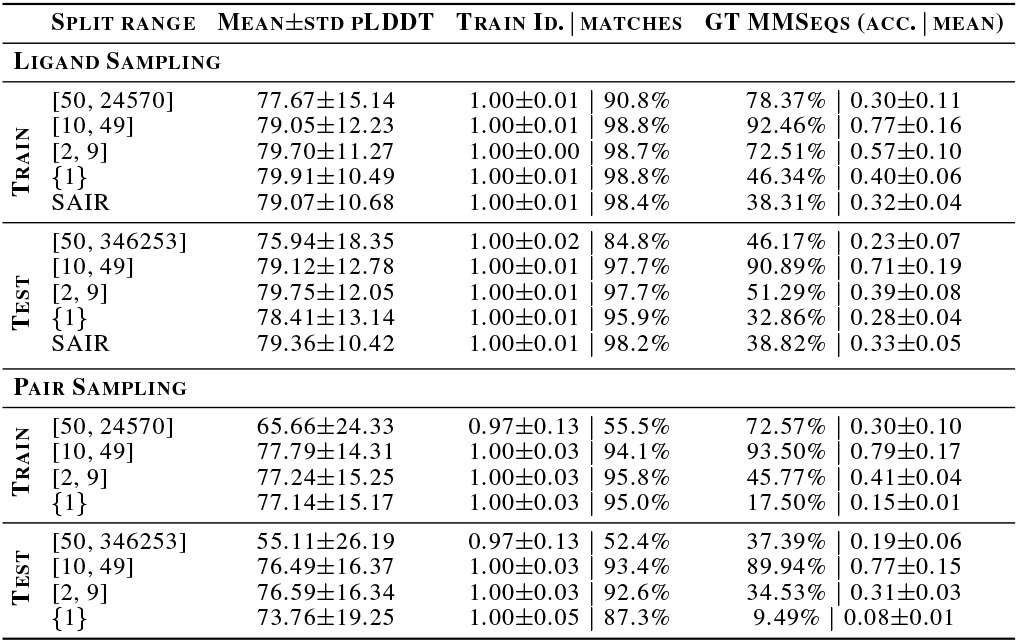
Per-stratum protein novelty statistics (MMSeqs2 similarity to the training set) together with pLDDT and GT retrieval. “Train Id. | matches” reports mean*±*std identity over generations with a valid MMSeqs hit, and the percentage of generations with any hit.

**Table A.3.** Per-input comparison of the model’s greedy decoding vs FP-NN under GT MMSeqs identity. FP-NN> indicates the fraction of inputs where FP-NN achieves higher maximum identity with respect to GT, and Model> the fraction where the model does. Δ_model>_ is the mean (model FP) identity among inputs where the model has higher identity. We additionally report the absolute FP-NN accuracy (mean best-match identity to GT across inputs) to contextualise the strength of retrieval under this metric.

| SPLIT RANGE | FP-NN ACC. | FP-NN $>$ | MODEL $>$ | TIES | MEAN $\Delta_{\text{MODEL}>}$ |
| --- | --- | --- | --- | --- | --- |
| [50, 346253] | 40.26% | 35.38% | 7.18% | 57.44% | 0.693 |
| [10, 49] | 46.62% | 28.50% | 22.50% | 49.00% | 0.890 |
| [2, 9] | 65.12% | 57.50% | 6.00% | 36.50% | 0.912 |
| {1} | 77.02% | 70.50% | 0.00% | 29.50% | – |

As a final note, these results should be interpreted with caution because GT MMSeqs rewards matching the specific annotated binder sequences and therefore favours retrieval mechanisms. Plausible binders from alternative families (or novel designs) are penalised even if they could bind, biasing towards retrieving the proteins present in the dataset.

### C. Extended–dataset results

Additionally, we also test results on the combination of the main binder dataset with the substrate–enzyme one. This combined dataset contains, for training, ∼1.8M unique ligands, ∼4.2M unique proteins and ∼13M total pairs, averaging 6.9 proteins per molecule. To avoid the most promiscuous ligands from overshadowing dataset statistics, the top 0.25% most common ones were removed, reducing the size to ∼1.8M unique ligands, ∼61K unique proteins and ∼5M pairs, averaging 3.3 proteins per molecule. Table A.4 shows how results converge back to retrieval behaviour, achieving largely similar scores to the Binder–Dataset.

### D. Maximally distant test set construction

We first run an all-to-all MMseqs2 search over dataset proteins and define an *effective identity* between proteins as 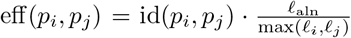, where id ∈ [0, 1] is the MMseqs2 identity, *ℓ*_aln_ the alignment length, and *ℓ*_*i*_, *ℓ*_*j*_ the protein lengths. Then, two proteins are considered connected if eff_*i,j*_ ≥ *τ*, with threshold *τ* = 0.85. When a protein is included in the test set, all connected proteins are also included, along with all their associated molecules. If the molecules are associated to other proteins, these are also included, causing a cascading effect. The algorithm ranks molecules by an estimated novelty score (average identity to the dataset scaled by number of connected neighbours) and iteratively adds connected sets. Cascades are killed when surpassing target test size, with examples at the boundary of the cascade not included, in order to prevent training leak. MMseqs2 search was performed with minimum coverage of 0.7, minimum alignment length of 50 and minimum sequence identity of 0.85. Using the aforementioned algorithm, a test set of 18,665 molecules was built. Additionally, we also created a test set of 400 molecules extracted from the SAIR dataset, which provides residue contact annotation, allowing to differentiate the binding pocket. For this set we do not maximise distance to training, instead, we simply ensure that its molecules are not present during training.

**Table A.4.** Results on Extended–Dataset: Main and substrate–enzyme. We report pLDDT, MMSeqs identity with percentage of matched proteins, and GT MMSeqs performance. SAIR test set is used instead of range-based ones (which maximised distance to train and were not as representative).

|  | SPLIT RANGE | PLDDT | TRAIN ID. | MATCHES | GT ACC. | MEAN |
| --- | --- | --- | --- | --- | --- | --- |
| TRAIN | [50, 134] | 77.98±14.72 | 1.00±0.02 | 89.9% | 79.52% | 0.32±0.13 |
|  | [10, 49] | 79.25±11.71 | 1.00±0.01 | 97.8% | 90.84% | 0.78±0.14 |
|  | [2, 9] | 79.65±10.92 | 1.00±0.01 | 98.5% | 65.98% | 0.54±0.10 |
|  | {1} | 78.77±11.30 | 1.00±0.00 | 97.9% | 48.34% | 0.44±0.04 |
|  | SAIR-400 | 79.35±10.33 | 0.99±0.00 | 98.7% | 47.00% | 0.41±0.06 |
| TEST | SAIR-400 | 70.62±22.78 | 0.99±0.02 | 66.6% | 31.24% | 0.27±0.03 |

**Figure A.2.**
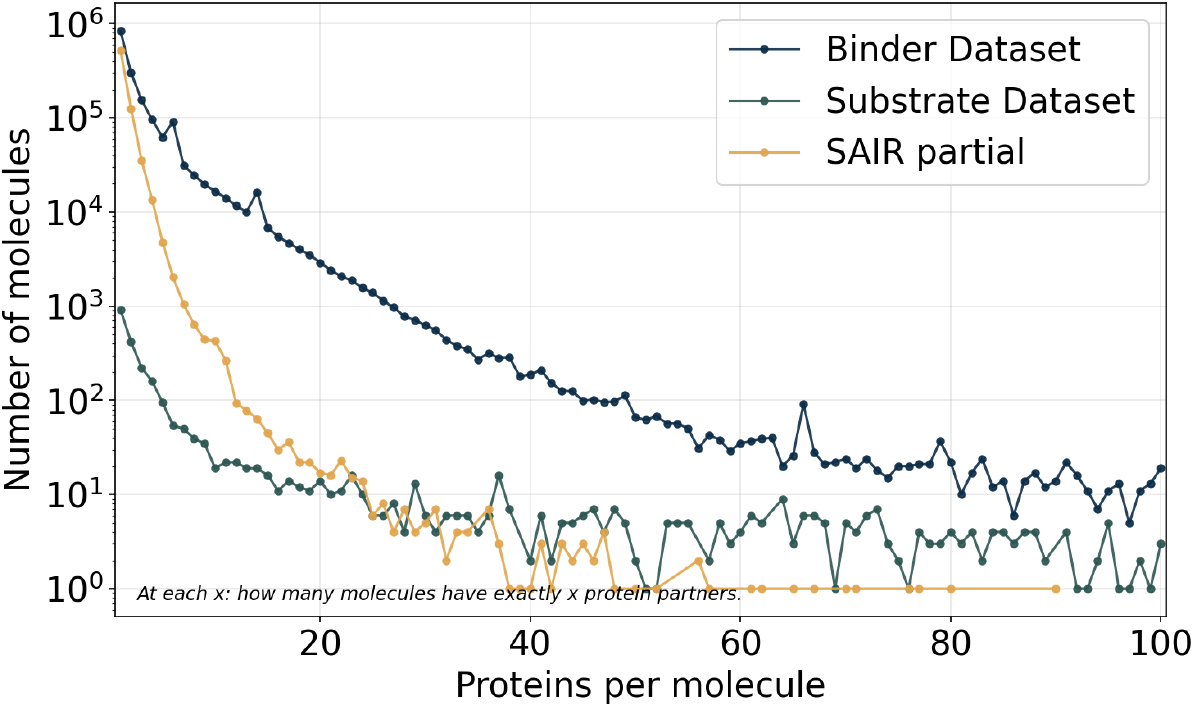
Distribution of protein partners per molecule. For each value of proteins per molecule (x-axis), the y-axis shows how many molecules have exactly that many unique protein partners. Curves are shown for the Binder Dataset, Substrate Dataset, and SAIR partial (log scale; x capped at 100).

### E. Pocket-only model

As a proof of concept, we also train a model to generate only the pocket region.

For this we leverage BioLip’s residue contact annotation. Using the same algorithm employed for SAIR, we estimate pocket sub-sequences in the amino acid chains, constructing a database of 34K unique ligands, 142K unique pocket sequences, and a total of 252K pairs. We deem appropriate to perform our experiment with this set rather than SAIR one, given that the average GT per ligand is ∼5, as opposed to SAIR’s ∼1. Notice that this database is orders of magnitude smaller than the Binder–dataset, given the scarcity of contact / pocket annotation. Consequently, the aim is not to achieve competitive performance, but to evaluate the learnability of the problem.

Results in Table A.5 show an average pLDDT for [2, 9] and {1} almost achieving the 70 cut-off, which is often used as reference for confident folding, and lower values for the more promiscuous strata. Regarding GT accuracy, it is substantially high for the least annotated ones in training, but falls in the test set, a sign of overfitting, most likely due to dataset size. Still, from this we extract that it is possible to associate the pocket sequences to the ligands they bind, but becomes harder when multiple pockets can accommodate that same ligand, as memorisation is less straight forward. With the current evidence we cannot claim whether this approach will still overfit to training with a larger database or, on the contrary, will become comparable to full protein approaches.

**Table A_5.**
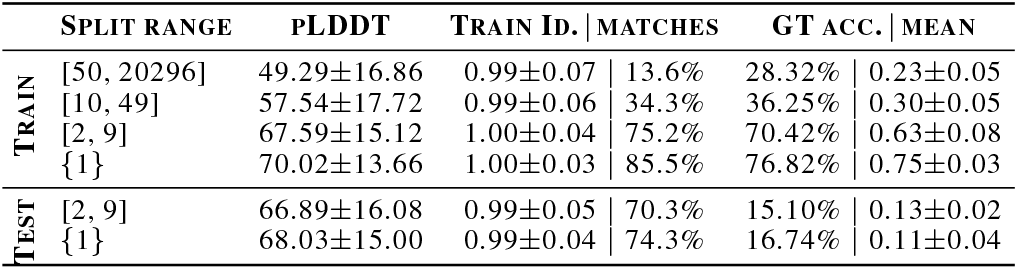
Results for pocket–only training.

## Footnotes

1 https://huggingface.co/collections/AI4PD/mol2pro-family

2 https://github.com/AI4PDLab/Mol2Pro

